# Mandatory use of mock communities highlighted by the descriptive comparison of Epi2Me 16S and EMU bioinformatic workflows for full-length 16S rRNA Nanopore sequencing

**DOI:** 10.64898/2026.08.04.742759

**Authors:** Fanie Shedleur-Bourguignon, William P. Thériault, Alexandre Thibodeau

**Affiliations:** Chaire de recherche en salubrité des viandes de l’Université de Montréal, Université de Montréal, Québec, Canada; Centre de recherche en infectiologie porcine et avicole (CRIPA)

## Abstract

Full-length 16S rRNA gene sequencing using Oxford Nanopore Technologies has emerged as a promising approach to improve species-level resolution in microbiota studies. However, the accuracy of taxonomic assignment remains highly dependent on the bioinformatics s used to process Nanopore long-read data. Therefore, the only way to ensure a good level of certainty in obtained results is to use positive controls in the form of mock communities in the experimental designs. In this study, we compared the performance of Epi2Me 16S (using Minimap2 or Kraken2) workflows provided by Oxford Nanopore Technologies and an EMU workflow for full-length 16S rRNA gene analysis. Using a commercial mock community sequenced across multiple Nanopore runs, taxonomic assignment accuracy and reproducibility was evaluated. Epi2Me-Kraken2 exhibited 18 % of incorrect genus-level assignments and failed to identify 3 species present in the mock community. While Epi2Me-Minimap2 achieved an excellent genus-level classification, reporting 9 % of sequences assigned to a genus not in the mock community, species-level assignments were inconsistent for several community members such as *Listeria*. In contrast, EMU provided accurate and consistent species-level taxonomic profiles, with all species correctly identified while keeping the number of genus absent from the mock community at 1.2%.

**Importance:** These results highlight that Epi2Me integrated workflows are not the best option for specie-level taxonomic assignation. More importantly, this paper underscores the importance of routine inclusion of positive controls for microbiota studies, in the form of mock communities, as a critical safeguard for accurate data interpretation. Without the use of a mock community, a paper published would be at risk of reporting wrong observations and inaccurate conclusions.

## Introduction

High-throughput sequencing has fundamentally transformed microbiology, ushering the field into the era of microbiome science. This advance has enabled a broad range of research avenues, from targeted manipulation of microbial communities for host health improvement to the investigation of complex, multiscale interactions such as the gut–brain axis. Concurrently, continuous technological improvements have reduced sequencing costs and increased accessibility, facilitating the adoption of microbiome analyses across research disciplines (1).

Within microbiota research, 16S rRNA gene amplicon sequencing remains the most widely used approach, owing to its cost-efficiency, scalability, and methodological robustness. The 16S rRNA gene is universally conserved among bacteria and archaea, while containing interspersed hypervariable regions that provide taxonomically informative sequence diversity (1). These variable regions are flanked by conserved sequences that enable the design of universal PCR primers, enabling amplification of defined variable regions suitable for taxonomic profiling. Among these, the V4 region is one of the most targeted, as it offers a favorable compromise between phylogenetic resolution and compatibility with short-read sequencing technologies (2).

In conventional short-read 16S rRNA amplicon sequencing, a selected hypervariable region is amplified from total extracted DNA, sample-specific barcodes are incorporated during library preparation, and libraries are subsequently pooled for sequencing, most commonly on Illumina platforms. Following data acquisition, reads undergo a bioinformatic processing that includes quality check and adjustments, clustering and taxonomic assignment base on comparison with reference databases. Sequences may be clustered into operational taxonomic units (OTUs), historically used as proxies for bacterial species, or resolved as exact amplicon sequence variants (ASVs). Although genus-level classification is reliable, species-level assignment remains challenging due to the limited phylogenetic information contained within short variable regions (3).

The emergence of long-read sequencing technologies, most notably developed by Oxford Nanopore Technologies, has enabled sequencing of full-length 16S rRNA sequences at a reasonable price. This technological shift substantially increases the potential for species-level taxonomic resolution in microbiota studies and represents a critical step toward overcoming the inherent limitations of short-read 16S amplicon sequencing. Recent literature found this technology offer clear advantage compared to traditional short amplicon 16S sequencing, allowing to identify sequences at the species-level and thus better biomarker discovery possibilities (4, 5).

However, this technology progress has created a gap for the analysis of the output raw reads. Most currently available 16S bioinformatic tools were originally designed and optimized for short read data (6). Nanopore sequencing is still more prone to introducing errors in the reads, compared to classic Illumina sequencing, even with the new chemistry (7, 8). These characteristics complicate downstream processing highlighting the need for analytical approaches specifically adapted to long-read error prone sequencing data.

Oxford Nanopore Technologies has developed an easy-to-use Windows-based platform, Epi2Me, which includes dedicated workflows for full-length 16S rRNA gene analysis, based on either Kraken2 or Minimap2 (9), providing user with a ready to use convivial solution. In parallel, several standalone bioinformatics tools specifically designed for full-length 16S Nanopore sequencing have emerged, including EMU (10), SituSeq (11), NaNoCLUST (12), and RESCUE (13) to name a few. These tools generally require non-Windows computational environments and are more demanding in terms of computational resources and skills. Each tools uses different ways to address the complexity of full length 16S Nanopore reads, details that are fully described in their respective publications. To date, none of these approaches has clearly emerged as a consensus or standard solution within the scientific community and currently the only way to ensure that these tools fits laboratory needs is by testing them, using well characterized sequencing data so that errors or strange outputs can be easily flagged.

In this context, the objective of the present study was to compare the full-length 16S workflows provided by Epi2Me (v1.3.0) with an in-house analysis workflow based on EMU (v3.5.1 and v3.6.2). At first, a single run of Nanopore sequencing was done and the sample containing the mock community was used for workflow comparisons. Then, as the sequencer started to be readily used in our research team for microbiota related projects, mock community samples were pull out of these runs (additional 5 independent runs) to assess the stability of the selected workflow, EMU. It is important to stress out that if projects did not run mock communities, our research team could have never detected that the workflows and options used needed some adjustments to output reliable results.

## Results

### Epi2Me-Minimap2

The performance of Epi2Me 16S-Minimap2 was evaluated while setting different parameters options (Table 1). At the genus level, it provided great classifications across parameter tested. However, species-level taxonomic assignment was unsatisfactory as for mock community members, such as *Listeria*, where species assignation was entirely wrong. The notable exceptions were *Salmonella enterica* and *Enterococcus faecalis*.

**Table 1:** Relative abundances of ZymoBIOMICS microbial community standard DNA according to different analysis options in Epi2Me using Minimap2. ()= number of falsely classification. For the specie taxonomic option, the best option is highlighted by ***. For the genus taxonomic assignation, the best option is highlighted by **. The lowest number of unclassified sequences is highlighted by *. The lowest number of wrongly classified genus (not supposed to be present in the mock community) is highlighted by ****. All analyses were made on the Mock 1 sample. Theo: theorical composition. COV: coverage option. SIM: similarity option. L.mono.: *Listeria monocytogenes*. P. aeru : *Pseudomonas aeruginosa*. Pseudo: *Pseudomonas*. Escher : *Escherichia*. S.ent : *Salmonella enterica*. Salmo: *Salmonella*. L.fermen : *Limosilactobacillus fermentum*. Limo: *Limosilactobacillus*. E. fae: *Enterococcus faecalis*. Enteroc: *Enterococcus*. Staph: *Staphylococcus*. Unclass: Unclassified. Bad genus: other species from genus absent from the theorical composition of the mock community.

| Bacteria | Theo | 95sim<br>90cov | 90sim<br>90cov | 90sim<br>95cov | 95sim<br>85cov |
| --- | --- | --- | --- | --- | --- |
| L.mono. | 0.141 | NA | <0.001 | <0.001 | <0.001 |
| Other Listeria | 0 | 0.075<br>(12) | 0.117<br>(15) | 0.112<br>(14) | 0.071<br>(12) |
| Total Listeria | 0.141 | 0.075 | 0.117** | 0.112 | 0.071 |
| P. aeru. | 0.042 | 0.013 | 0.029*** | 0.017 | 0.014 |
| Other Pseudo | 0 | <0.001<br>(1) | 0.001<br>(14) | <0.001<br>(13) | <0.001<br>(1) |
| Total Pseudo | 0.042 | 0.013 | 0.029** | 0.017 | 0.014 |
| B. subtilis | 0.174 | 0.017 | 0.024*** | 0.024*** | 0.015 |
| Other Bacillus | 0 | 0.143<br>(13) | 0.222<br>(44) | 0.214<br>(43) | 0.140<br>(19) |
| Total Bacillus | 0.174 | 0.160** | 0.246 | 0.238 | 0.155 |
| E. coli | 0.101 | <0.001 | <0.001 | <0.001 | <0.001 |
| Other Escher | 0 | 0.018<br>(3) | 0.04<br>(3) | 0.037<br>(3) | 0.022<br>(3) |
| Total Escher | 0.101 | 0.018 | 0.040** | 0.037 | 0.022 |
| S.ent | 0.104 | 0.067 | 0.109*** | 0.085 | 0.073 |
| Other Salmo | 0 | 0.002 | 0.002 | 0.002 | 0.002 |

Table 1: Relative abundances of ZymoBIOMICS microbial community standard DNA according to different analysis options in Epi2Me using Minimap2.
|  |  | (1) | (1) | (1) | (1) |
| --- | --- | --- | --- | --- | --- |
| Total Salmo | 0.104 | 0.069 | 0.111** | 0.087 | 0.075 |
| L.fermen | 0.184 | 0.060 | 0.094*** | 0.072 | 0.052 |
| Other Limo | 0 | 0 | <0.001<br>(2) | <0.001<br>(2) | (0) |
| Total Limo | 0.184 | 0.060 | 0.094** | 0.072 | 0.052 |
| E. fae | 0.090 | 0.044 | 0.073*** | 0.068 | 0.043 |
| Other Enteroc | 0 | 0.001<br>(8) | 0.004<br>(19) | 0.003<br>(12) | 0.001<br>(8) |
| Total Enteroc | 0.090 | 0.045 | 0.077** | 0.071 | 0.044 |
| S. aureus | 0.155 | 0.066 | 0.080*** | 0.075 | 0.06 |
| Other Staph | 0 | 0.048<br>(13) | 0.088<br>(36) | 0.083<br>(37) | 0.045<br>(16) |
| Total Staph | 0.155 | 0.114 | 0.168 | 0.158** | 0.051 |
| Bad genus | 0 | 0.046****<br>(29) | 0.118<br>(118) | 0.208<br>(108) | 0.096<br>(37) |
| Unclass | 0 | 0.400 | 0.038* | 0.137 | 0.420 |

Increasing stringency by applying higher similarity (sim) and coverage (cov) thresholds reduced the number of incorrectly assigned genera but resulted in a substantial increase in unclassified reads. The most stringent parameter combination tested (95% sequence similarity and 90% alignment coverage) minimized false genus-level assignments but also produced the highest proportion of unclassified sequences, highlighting a trade-off between classification accuracy and sensitivity. The gain for correct species assignment was marginal when varying parameters.

### Comparison of Epi2Me and EMU

The best-performing Epi2Me-Minimap2 configuration (90 cov, 90 sim), the Epi2Me-Kraken2 (default parameters) and the custom EMU workflows were subsequently compared using the same mock community sample sequenced on the same flowcell (same run) (Table 2, Figure 1 and Figure 2). At the genus level (Table 1 and Figure 1), results are comparable between all workflows except for Kraken2 that exhibited poor overall performance, characterized by a high number and high relative abundance of incorrectly assigned genera: in total, more than 200 genus-level misclassifications were observed.

**Figure 1:**
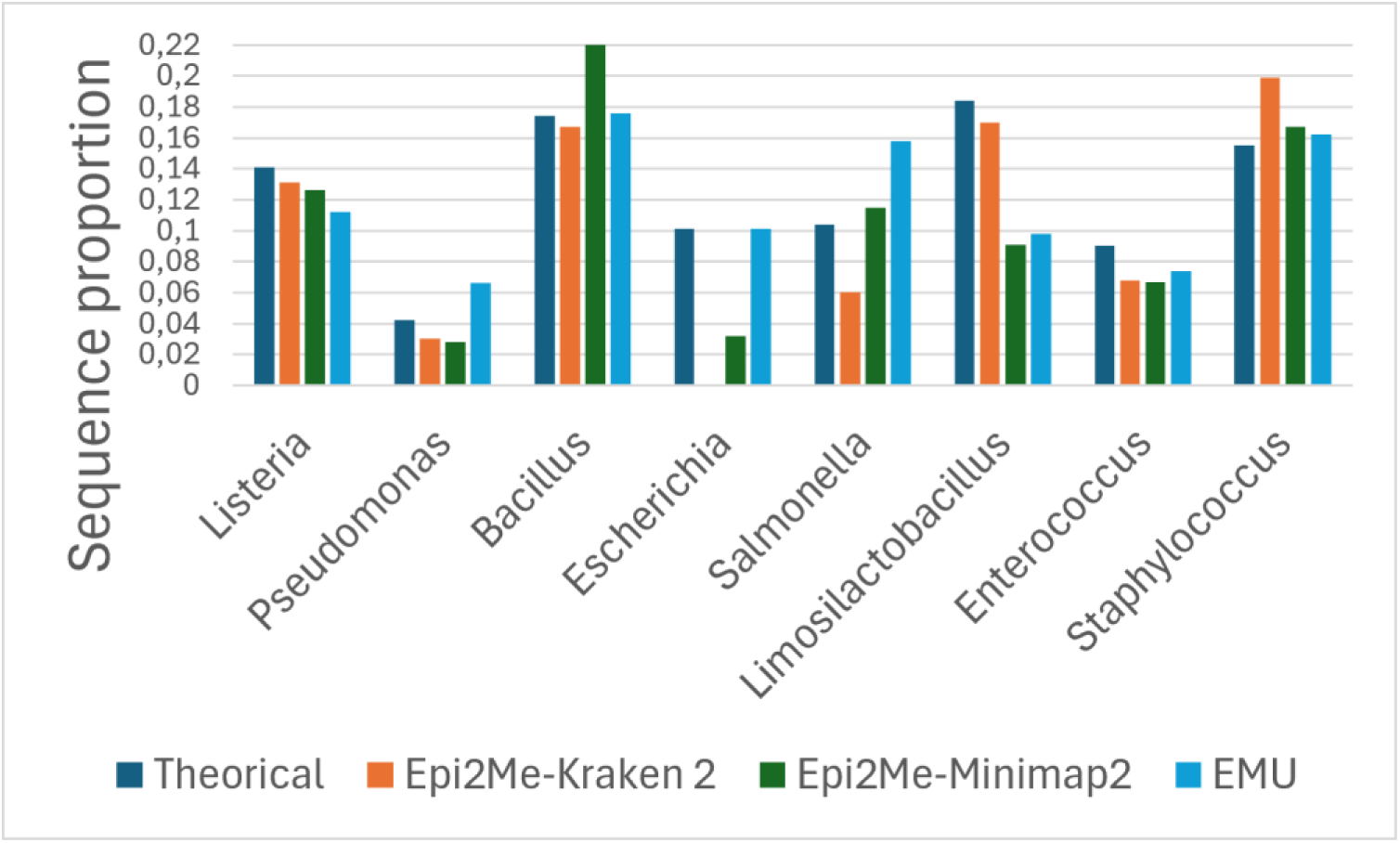
Comparison of Epi2Me-Kraken2, Epi2Me-Minimap2 and EMU workflows for taxonomic assignation at the genus level, in relative abundance.

**Figure 2:**
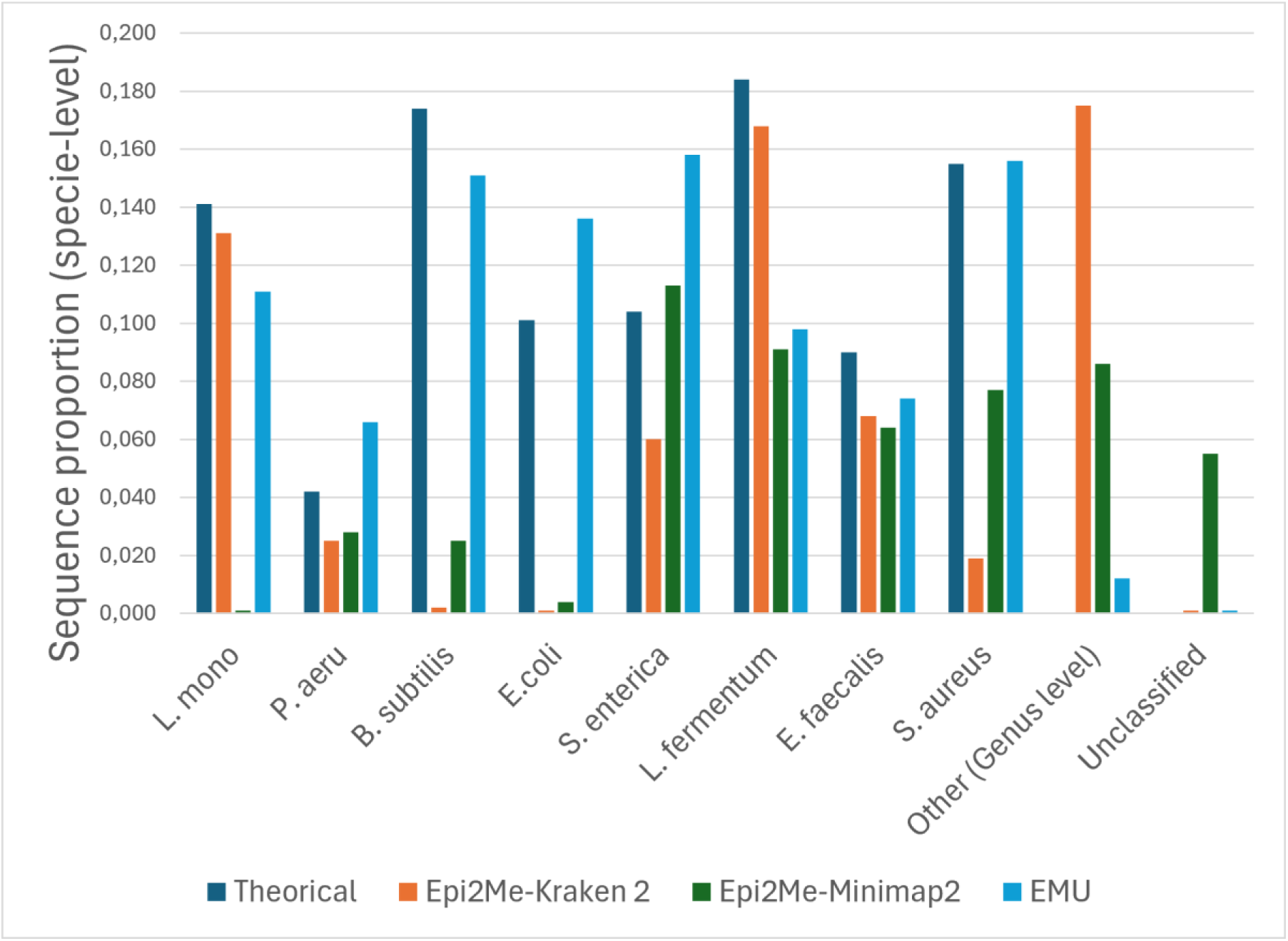
Comparison of Epi2Me-Kraken2, Epi2Me-Minimap2 and the EMU workflows for taxonomic assignation at the species-level, in relative abundances. Other (Genus level): species classified as genus absent from the theorical composition of the mock community.

**Table 2:** Comparison of Epi2Me-Kraken2, Epi2Me-Minimap2 and the custom EMU workflow for taxonomic assignation at the specie and genus level in relative abundances. ()= number of falsely classification. For the specie taxonomic option, the best option is highlighted by ***. For the genus taxonomic assignation, the best option is highlighted by **. The lowest number of unclassified sequences is highlighted by *. The lowest number of wrongly classified genus (not supposed to be present in the mock community) is highlighted by ****. All analyses were made on the Mock 1 sample. Theo: theorical composition. COV: coverage option. SIM: similarity option. L.mono.: *Listeria monocytogenes*. P. aeru : *Pseudomonas aeruginosa*. Pseudo: *Pseudomonas*. Escher : *Escherichia*. S.ent : *Salmonella enterica*. Salmo: *Salmonella*. L.fermen : *Limosilactobacillus fermentum*. Limo : *Limosilactobacillus*. E. fae: *Enterococcus faecalis*. Enteroc: *Enterococcus*. Staph: *Staphylococcus*. Unclass: Unclassified. Bad genus: other species from genus absent from the theorical composition of the mock community.

| Bacteria | Theo (%) | Kraken2 default | Epi2me Best | EMU |
| --- | --- | --- | --- | --- |
| L.mono. | 0.141 | 0.131*** | <0.001 | 0.111 |
| Other Listeria | 0 | 0.001 (9) | 0.126 (19) | 0.002 (4) |
| Total Listeria | 0.141 | 0.131** | 0.126 | 0.112 |
| P. aeru. | 0.042 | 0.025 | 0.028 | 0.066*** |
| Other Pseudo | 0 | 0.005 (50) | <0.001 (9) | 0 (0) |
| Total Pseudo | 0.042 | 0.030 | 0.028 | 0.066** |
| B. subtilis | 0.174 | 0.002 | 0.025 | 0.151*** |
| Other Bacillus | 0 | 0.165 (63) | 0.208 (40) | 0.025 (13) |
| Total Bacillus | 0.174 | 0.167 | 0.233 | 0.176** |
| E. coli | 0.101 | <0.001 | 0.004 | 0.136*** |
| Other Escher | 0 | <0.001 (2) | 0.028 (3) | 0.001 (2) |
| Total Escher | 0.101 | <0.001 | 0.032 | 0.101** |
| S.ent | 0.104 | 0.060 | 0.113*** | 0.158 |
| Other Salmo | 0 | <0.001 (1) | 0.002 (1) | 0 |
| Total Salmo | 0.104 | 0.060 | 0.115** | 0.158 |
| L.fermen | 0.184 | 0.168*** | 0.091 | 0.098 |
| Other Limo | 0 | 0.002 | <0.001 | 0 |

Table 2: Comparison of Epi2Me-Kraken2, Epi2Me-Minimap2 and the custom EMU workflow for taxonomic assignation at the specie and genus level in relative abundances.
|  |  | (11) | (3) |  |
| --- | --- | --- | --- | --- |
| Total Limo | 0.184 | 0.170** | 0.091 | 0.098 |
| E. fae | 0.090 | 0.068 | 0.064 | 0.074*** |
| Other Enteroc | 0 | <0.001<br>( ) | 0.003<br>(33) | <0.001<br>(2) |
| Total Enteroc | 0.090 | 0.068 | 0.067 | 0.074** |
| S. aureus | 0.155 | 0.019 | 0.077 | 0.156*** |
| Other Staph | 0 | 0.180<br>(32) | 0.090<br>(42) | 0.008<br>(16) |
| Total Staph | 0.155 | 0.199 | 0.167** | 0.162 |
| Bad genus | 0 | 0.175<br>(>200) | 0.086<br>(50) | 0.012****<br>(17) |
| Unclass | 0 | <0.001* | 0.055 | <0.001* |

At the species-level, compared with Epi2Me-Minimap2, EMU workflow provided improved taxonomic assignments (Table 1, Figure 2). Several species that were poorly detected or misclassified by Epie2Me-Minimap2, including *Listeria monocytogenes*, *Bacillus subtilis*, and *Escherichia coli*, were accurately identified using EMU. In addition, it consistently yielded fewer incorrectly assigned genera, both in terms of absolute numbers and sequence relative abundances. Nevertheless, none of the evaluated workflow achieved perfect classification across all mock community members, a result that was expected.

### Reproducibility of EMU workflow across sequencing runs

The output classification stability using the custom EMU workflow was assessed across six independent Nanopore sequencing runs (Table 3, Figure 3). Although some variability in relative abundance estimates was observed between runs, the overall taxonomic profiles of the mock community remained largely consistent.

**Figure 3:**
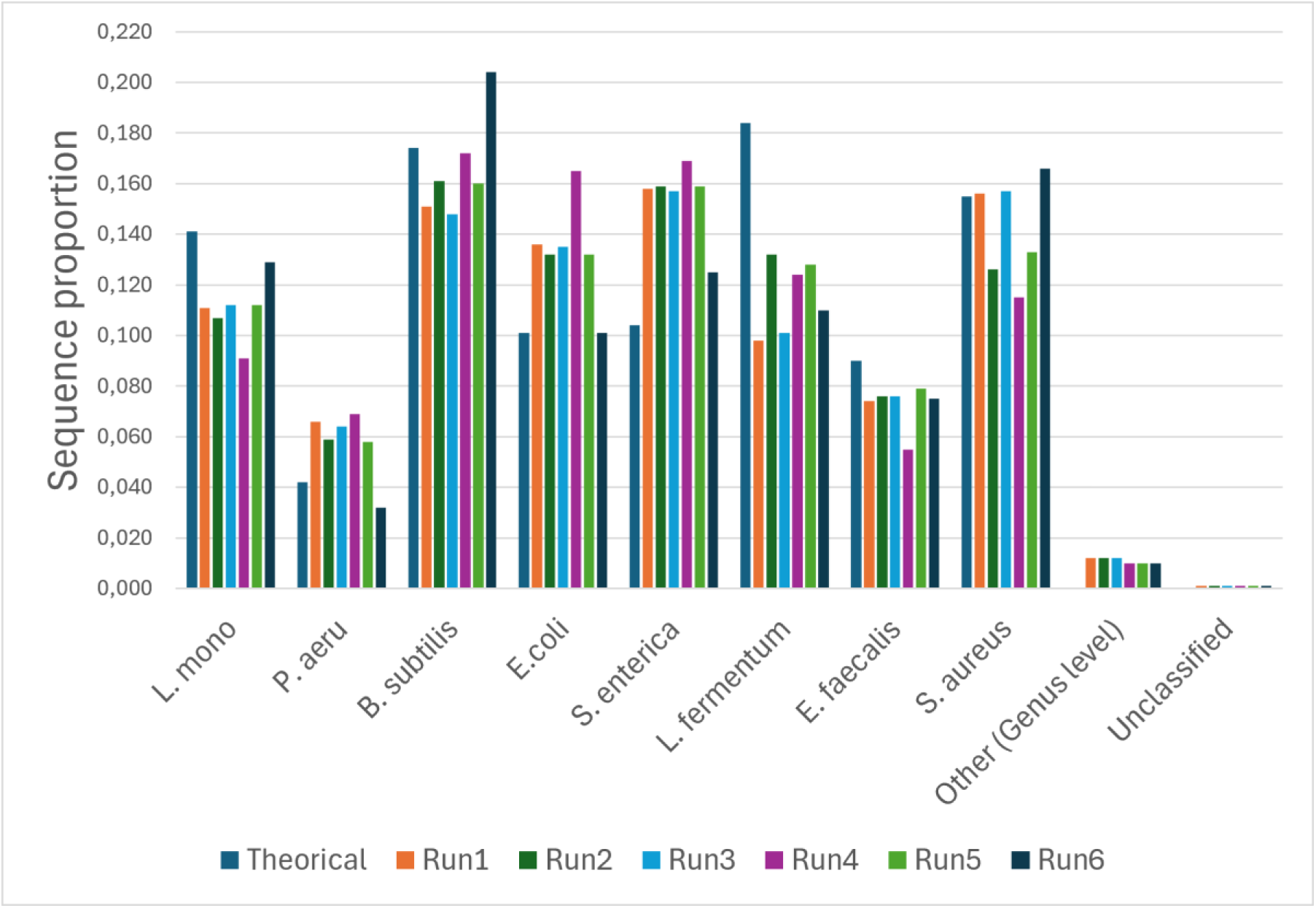
Multiple mock community samples comparison in relative abundances using EMU. Other (Genus level): species classified as genus absent from the theorical composition of the mock community.

**Table 3:** Multiple mock community samples comparison in relative abundances using EMU. ()= number of falsely classification. For the specie taxonomic option, the best option is highlighted by ***. For the genus taxonomic assignation, the best option is highlighted by **. The lowest number of unclassified sequences is highlighted by *. The lowest number of wrongly classified genus (not supposed to be present in the mock community) is highlighted by ****. All analyses were made on the Mock 1 sample. Theo: theorical composition. COV: coverage option. SIM: similarity option. L.mono.: *Listeria monocytogenes*. P. aeru : *Pseudomonas aeruginosa*. Pseudo: *Pseudomonas*. Escher : *Escherichia*. S.ent : *Salmonella enterica*. Salmo: *Salmonella*. L.fermen : *Limosilactobacillus fermentum*. Limo : *Limosilactobacillus*. E. fae: *Enterococcus faecalis*. Enteroc: *Enterococcus*. Staph: *Staphylococcus*. Unclass: Unclassified. Bad genus: other species from genus absent from the theorical composition of the mock community.

| Bacteria | Theo | Mock1 | Mock2 | Mock3 | Mock4 | Mock5 | Mock6 |
| --- | --- | --- | --- | --- | --- | --- | --- |
| L.mono. | 0.141 | 0.111 | 0.107 | 0.112 | 0.091 | 0.112 | 0.129 |
| Other Listeria | 0 | 0.002<br>(4) | 0.002<br>(4) | 0.002<br>(4) | 0.001<br>(4) | 0.001<br>(4) | 0.002<br>(4) |
| Total Listeria | 0.141 | 0.113 | 0.109 | 0.114 | 0.092 | 0.113 | 0.131 |
| P. aeru. | 0.042 | 0.066 | 0.059 | 0.064 | 0.069 | 0.058 | 0.032 |
| B. subtilis | 0.174 | 0.151 | 0.161 | 0.148 | 0.172 | 0.160 | 0.204 |
| Other Bacillus | 0 | 0.025<br>(15) | 0.024<br>(15) | 0.026<br>(15) | 0.021<br>(15) | 0.021<br>(15) | 0.029<br>(15) |
| Total Bacillus | 0.174 | 0.176 | 0.185 | 0.174 | 0.193 | 0.181 | 0.233 |
| E. coli | 0.101 | 0.136 | 0.132 | 0.135 | 0.165 | 0.132 | 0.101 |
| Other Escher | 0 | 0.001<br>(2) | 0.001<br>(2) | 0.001<br>(2) | 0.001<br>(2) | 0.001<br>(2) | 0.001<br>(2) |
| Total Escher | 0.101 | 0.137 | 0.133 | 0.136 | 0.166 | 0.133 | 0.102 |
| S.ent | 0.104 | 0.158 | 0.159 | 0.157 | 0.169 | 0.159 | 0.125 |
| L.fermen | 0.184 | 0.098 | 0.132 | 0.101 | 0.124 | 0.128 | 0.110 |
| E. fae | 0.090 | 0.074 | 0.076 | 0.076 | 0.055 | 0.079 | 0.075 |
| Other Enteroc | 0 | <0.001<br>(2) | <0.001<br>(2) | <0.001<br>(2) | <0.001<br>(2) | <0.001<br>(2) | <0.001<br>(2) |
| Total Enteroc | 0.090 | 0.074 | 0.076 | 0.076 | 0.055 | 0.079 | 0.075 |
| S. aureus | 0.155 | 0.156 | 0.126 | 0.157 | 0.115 | 0.133 | 0.166 |
| Other Staph | 0 | 0.008<br>(16) | 0.008<br>(16) | 0.008<br>(16) | 0.006<br>(16) | 0.005<br>(16) | 0.011<br>(16) |
| Total Staph | 0.155 | 0.162 | 0.134 | 0.165 | 0.121 | 0.138 | 0.177 |
| Bad genus | 0 | 0.012<br>(17) | 0.012<br>(17) | 0.012<br>(17) | 0.010<br>(17) | 0.010<br>(17) | 0.010<br>(17) |
| Unclass | 0 | <0.001 | <0.001 | <0.001 | <0.001 | 0.002 | <0.001 |

### General computational performance and EMU --N parameter variation

On the Digital Alliance of Canada server Narval, across the six mock community independent samples, evaluating raw versus processed reads (with chopper) and four different values of the --N parameter (1, 25, 50, and 100), resulting in a total of 48 EMU executions, required 19 hours of computer work using 25 CPU cores and the full 250 GB of allocated RAM. Analyses using unfiltered raw reads failed due to resource limitations, whereas runs using quality-trimmed reads (chopper) completed successfully.

Variation of the --N parameter influenced classification. Using --N of 1, 25, 50 and 100 gave respectively a total of 90 species, 68 species, 64 species and 72 species. For the same analysis, respectively, mean unclassified sequences, in relative abundance, were 0.003, 0.0008, 0.0007 and 0.0006. For the same analysis, the number of species classified as a genus other than the one theoretically in the mock community was 82, 60, 56 and 64.

### EMU version 3.6.2

While writing this manuscript, EMU was updated. EMU version 3.6.2, with the taxonomy database version March 2026, was therefore tested with the new option PID set at 0 (default), 50, 80 and 90. When using default parameters, the main difference of the new version, regarding the mock community, was that *Bacillus subtilis* was not found, but instead was *Bacillus spizizenii*. In the assignment distributions files outputted by EMU (--keep-read-assignments), the median probability assigned to *B. spizizenni* was of 0.64, while the median probability for *B. subtilis* was 0.09, illustrating that EMU is struggling to separate the two. For a long time, *B. spizizenii* was classified as a subspecie of *B. subtilis* (14) which might explain the different classification obtained. Some sequences downloaded into NCBI (used for EMU database) and annotated as *B. subtilis* might have been *B. spizienni*, sketching EMU database. The other possibility is that the strain used for the mock community was in fact a *B. spizienni* rather then a “true” *B. subtilis*. Eighter way, this misclassification seems more like a problem with the database nomenclature update following the introduction of *B. spizienni* then a problem with EMU per say.

Varying the PID also had effects on species assignation (Figure 4). The numbers of returned species for PID 0 (default), PID 80 and PID 90 were respectively: 85, 85 and 78. Notably, at PID 90, the number of unclassified sequences was the highest to 9 %, the number of sequences not supposed to be in the mock community increased to 16 % and the number or correctly classified *Bacillus bacilis*+*spizicenii* dropped to 3 %. AT PID 80, practically no change could be observed compared to default parameters.

**Figure 4:**
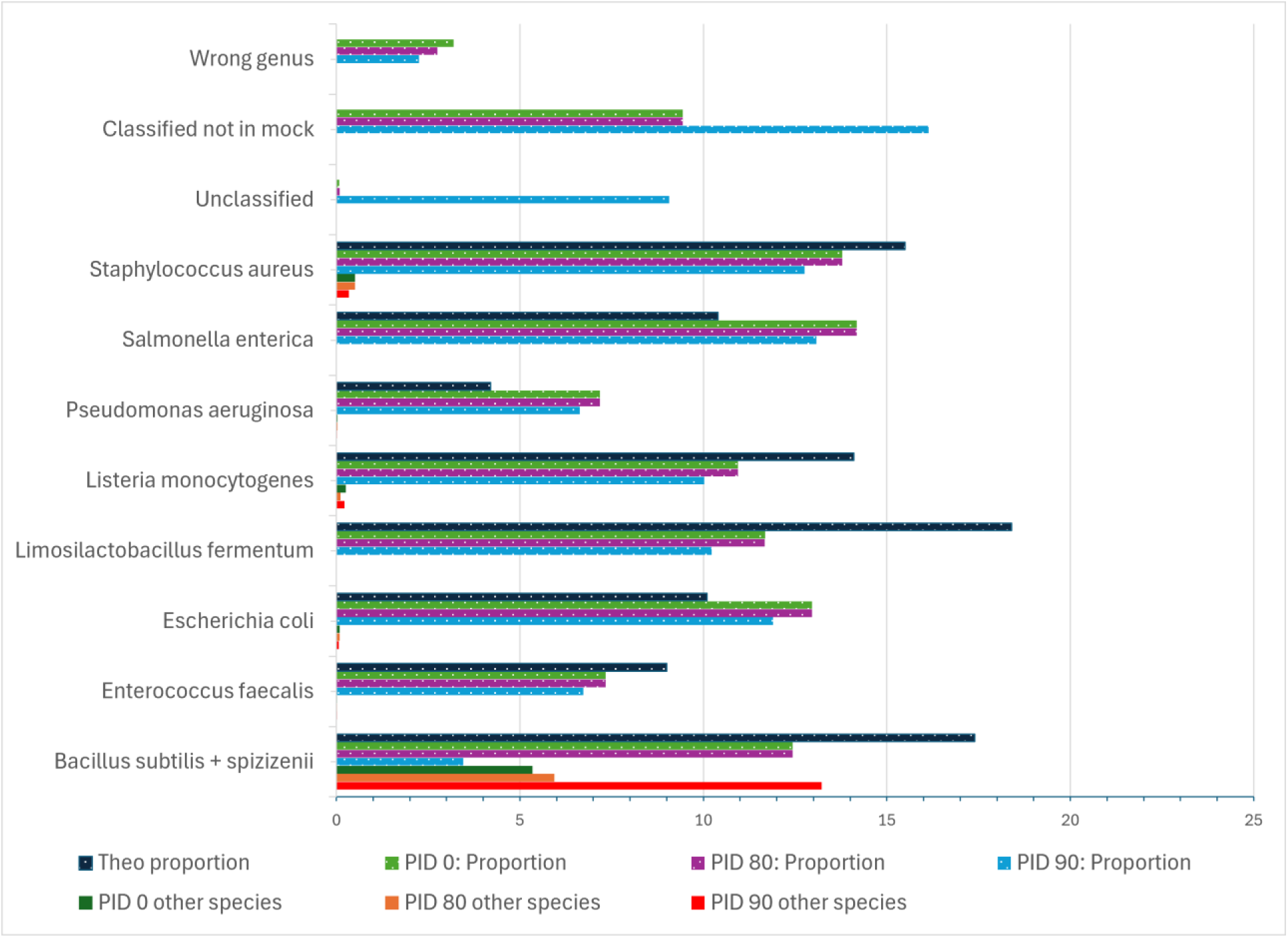
Proportions (%) of sequences taxonomic assignation when comparing different PID option values in EMU. Wrong genus: classified sequence at a genus not supposed to be in the mock community; other species: sequences classified at the correct genus but the incorrect species.

## Discussion

The observations reported in this study clearly highlight that using mock communities for microbiota sequencing studies is crucial as it is the only way to increase confidence in results and spot major errors that might originate from the bioinformatic analysis process. As new analytical tools and updated versions of existing software continue to emerge, systematic benchmarking using mock communities remains essential to rigorously assess methodological performance and robustness of selected workflows.

Among the bioinformatics approaches evaluated in this study, the custom EMU workflow provided the overall most accurate species-level taxonomic classification for full-length 16S rRNA gene using Nanopore sequencing. This was also observed in other studies (4, 9, 15)

Across the evaluated workflows, substantial differences in taxonomic assignation performance were observed. Epi2Me-Kraken2 exhibited notable limitations, particularly for several key mock community members, including *Bacillus subtilis*, *Escherichia coli*, and *Staphylococcus aureus*. Similarly, Epi2Me-Minimap2 demonstrated reduced accuracy to assign sequences to *Listeria monocytogenes*, *Bacillus subtilis*, and *Escherichia coli*, and showed reduced performance for *Staphylococcus aureus*. In contrast, the EMU yielded consistent and accurate classifications across species.

It should be noted, however, that the present study evaluation relied on a rather simple mock community added as DNA at the 16S PCR step. Further validation using more complex and compositionally diverse mock communities, including live communities that must be DNA extracted, is desirable. Regardless, underperformance of the Epi2Me relative to alternative bioinformatic tool was evident in our study and has been reported previously (12). Therefore, one must be careful with the interpretation of results generated through Epi2Me, especially if the authors forgot to use a mock community as control.

Others have tested EMU and some reported problems; for example, a marked underrepresentation of *Pseudomonas* (0.5% vs the expected 4%) was observed (9). However, such discrepancies were not observed in this study. This might be attributable to the use of an updated sequencing chemistry (R10 vs R9), combined with additional sequence cleaning steps prior to EMU. This also clearly demonstrate that comparison of different papers using significant different route to obtain sequencing data and species-level taxonomic assignation must be made with critical thoughts.

The near absence of unclassified reads in EMU outputs also warrants careful consideration. As documented in the EMU GitHub issue tracker, reads are reported as unclassified only when no alignment can be detected; sequences with even weak or potentially spurious alignments can be assigned a taxonomic label. This behavior could contribute to the low proportion of unclassified reads observed in this study as some low abundant species might be false-positive assignations. Although this characteristic raises concerns regarding potential “over-classification” or false positive species assignations, its practical impact appeared limited under the conditions evaluated here. Using the newest EMU version with the option PID did not modify the mock community in an impactful manner, up PID set at 80, and performances decreased at a stringent PID set at 90. Moreover, only a small proportion of reads were assigned to genera absent from the theoretical mock community composition, and these assignments accounted for a small fraction of total reads. Notable exceptions were observed for *Bacillus* and *Staphylococcus*, for which misassigned species represented up to 2% of each genus reads. This proportion is not negligible and warrants consideration when interpreting species-level profiles.

Disentangling the sources of taxonomic misclassification remains challenging. Errors may stem from limitations inherent to bioinformatic workflows, inaccuracies or incompleteness within reference databases, or residual sequencing errors associated with Nanopore technology (6). Some apparent errors also might not be real errors: as taxonomy evolves, new species are introduced or species are renamed, which may confuse users as database being updated with the latest taxonomy. It is likely that misclassification reflects a combination of methodological and technological factors rather than a single predominant source. Contributions from near unavoidable low-level contamination during sample processing or from reagents cannot be excluded (16), therefore meaning that a certain level of identification of genus absent from the theorical mock community composition is unavoidable and thus a negative control could be included to evaluate this. This control is run in our lab but not presented in the manuscript to keep the article focus on mock communities used as positive controls. Furthermore, excessive sequencing depth may further exacerbate the detection of spurious low-abundance taxa, amplifying errors to detectable levels (16, 17); false signals if enough abundant may then be interpreted as genuine species-level assignments by bioinformatic tools.

Given the rapid evolution of Nanopore sequencing chemistry, continuous protocol optimization remains critical. To minimize erroneous taxonomic assignments, users should aim to reduce the number of PCR amplification and barcoding cycles to the minimum required and to use high-quality, non-degraded DNA as input material. The lowest error-prone TAQ polymerase, available to researcher, being able to be used is also a consideration. Optimizing sequencing depth according to sample complexity may further mitigate the impact of over-sequencing. However, the influence of wet-lab protocol parameters on error profiles is likely sample-dependent and remains insufficiently characterized. This gap is reflected in the limited number of studies addressing the influence of laboratory (DNA extraction to library preparation) and sequencing protocol variations on taxonomic profiles in full 16S rRNA gene analyses (3, 6, 18).

Overall, observation of the present study supports the growing body of literature demonstrating that full-length 16S rRNA gene sequencing using Nanopore technology represents a viable approach for microbiota profiling, provided that its methodological limitations are explicitly acknowledged and taken into consideration by authors when writing (7). Species-level taxonomic assignments, particularly for low-abundance taxa, should be interpreted with caution, especially when central to study conclusions. As always, important discoveries should be validated using complementary methodologies. Diversity analyses at the species-level remain informative despite a degree of taxonomic uncertainty, as such analyses rely primarily on sequences being “different” rather than exact taxonomic identities.

More importantly, the observation of this work adds to a call made by members of the microbiota community to increase reproducibility in sequencing studies (19). The authors hope that it helps researchers to reach the clear conclusion that the routine inclusion of mock communities sequenced in parallel with experimental samples is mandatory to assess run-to-run variability and contextualize classification errors. Mock communities should also be used even when using well established bioinformatic tools as new database of software versions might cause changes. Also, all researchers are at risk of inputting coding errors in their favorite workflow, errors that might change the results without the user notifying if no controls are used. As full-length 16S sequencing becomes increasingly accessible, continued methodological refinement and transparent reporting will be essential to ensure robust reproducible and credible microbiome research. Careful conclusions in microbiota studies should be made by having a deep look at the data and never blindly trust bioinformatic workflow results.

## Materials and Methods

### Library preparation and sequencing

The ZymoBIOMICS Microbial DNA Standard (Zymo Research, California, USA) was used as the mock community for full-length 16S rRNA gene sequencing. Full-length 16S rRNA gene amplification was performed using primers NANO_27_F (TTTCTGTTGGTGCTGATATTGCAGAGTTTGATYMTGGCTCAG) and NANO_1492_R (ACTTGCCTGTCGCTCTATCTTCTACGGYTACCTTGTTACGACTT). PCR reactions were carried out using Invitrogen Platinum SuperFi Taq (ThermoFisher, Ontario, Canada), 0.2mM dNTPs, 0.6 µM of each primer, BSA 0.4mg/ml and 25 ng of DNA for each sample. The cycling conditions were: 5 minutes at 95°C for polymerase activation followed by cycles of 30 s at 95°C, 30 s at 55°C, 3 min at 72°C and the PCR ended with a final extension of 10 min at 72 °C. A total of 23 amplification cycles was applied.

Amplicons were subsequently barcoded using the Oxford Nanopore PCR Barcoding Expansion kit EXP-PBC096 (Nanopore, Oxford, United-Kingdom) according to the manufacturer’s instructions. PCR reactions were carried out using Invitrogen Platinum SuperFi Taq (ThermoFisher), 0.2mM dNTPs, 0.5 µl of barcode and 200 ng of PCR amplicon. The following cycling conditions were used: 3 min at 95°C for polymerase activation followed by cycles of 15 s at 95°C, 15 s at 62°C, 2 min at 72°C and ending with a final extension of 10 min at 72 °C. A total of 15 amplification cycles was applied.

Barcoded amplicons were pooled into a single library and purified using Beckman Coulter Canada Lp AMPure XP (ThermoFisher) following the recommended protocol. Adapter ligation was performed using the Oxford Nanopore Ligation Sequencing Kit v14 (Nanopore) as per requirements by the manufacturer. The final library was loaded onto an R10.4.1 flow cell (Nanopore) and sequenced on a Minion (Nanopore) for 72 hours to maximize pore utilization as per the manufacturer requirements.

### Raw sequence processing and taxonomic analysis

Basecalling was done in parallel of sequencing using Guppy (v6.5.7), with the super-accurate basecalling model, on an RTX 3070 GPU. Adapter trimming was enabled, reads containing internal adapter sequences were discarded, and samples were demultiplexed during basecalling.

For taxonomic profiling using the Epi2Me 16S workflows (v1.3.0), two classification strategies were evaluated: Kraken2 and Minimap2. The default NCBI reference database provided by Epi2Me was used for both approaches. Kraken2 and Minimap2 analyses were first performed using default parameters. For Minimap2, as this is also the base of EMU, multiple combinations of sequence similarity and alignment coverage thresholds were tested, as detailed in the results tables, to try to maximise sequences species assignment accuracy. Resulting taxonomic abundance tables were exported and analyzed in Microsoft Excel.

For EMU workflow, demultiplexed FastQ files corresponding to each barcode were concatenated into a single file per sample using the Windows command prompt (copy /b *.fastq.gz sample_barcode.fastq.gz). Sequencing quality was assessed using FastQC (v0.12.1), with particular attention to quality score variation at the 5′ and 3′ ends of reads.

Because sequencing quality profiles varied between Nanopore runs, read trimming was performed on a per-run basis. Reads were trimmed using Chopper (v0.10.0b) to remove low-quality nucleotides from the beginning (headcrop) and/or end (tailcrop) until a mean quality score graph of Q20 was achieved; this corresponded to a 20-30 nucleotides removal at each end. After run-based trimming, all samples were processed in the same way, regardless of the sequencing run. Chopper was used to retain reads within a length range of 1300–1700 bp (--minlength 1300; --maxlength 1700) and with a minimum quality score of 15 (--quality 15). To adjust for sequencing depth, Seqtk (v1.5-r133) was used to subsample each dataset to a maximum of 150,000 reads. This was required as some mock communities were sequenced in runs where very few samples were loaded.

Processed reads were analyzed using EMU (v3.5.1), with the minimum relative abundance threshold set to 0.000001 (--min-abundance 0.000001) and the other options used with default values, unless otherwise specified. The default EMU NCBI database was downloaded (V3.4.5) and used for taxonomic assignment. EMU outputs were combined at the species-level using the combine-outputs function. Resulting tables were examined using Microsoft Excel. Complemental analyses were conducted to evaluate the impact of using unfiltered raw reads (no Chopper) and varying the --N parameter (1, 25, 50, and 100) on taxonomic classification accuracy.

As this article was in the final steps of production, new EMU (3.6.2) and taxonomic database (March 2026) versions became available. EMU 3.6.2 offers the possibility to set further options such as minimum percent identity (PID), based on NM tag and aligned query length requirements for read filtering before taxonomic assignation, that has the potential to reduce false positive species assignation. The trimmed mock community samples were therefore also analyzed by these new versions (EMU and database) with different PID options: 0 (default), 80 and 90.

On a side note, read concatenation and FastQC analyses were performed on a desktop PC, whereas Chopper and EMU were executed on the Digital Research Alliance of Canada Narval cluster using 32 CPU cores and 250 GB of RAM. Output files were then downloaded back to a desktop PC for analysis.

### AI-assisted language editing statement

The manuscript underwent language editing with the assistance of an AI-based large language model (ChatGPT, OpenAI) after the first manuscript draft. The tool was used exclusively to improve the clarity, coherence, and academic style of the English language, including grammar, syntax, and phrasing, and to harmonize the writing across sections. No scientific content, data, results, interpretations, or conclusions were generated, altered, or influenced by the AI. All methodological choices, analyses, and interpretations remain the sole responsibility of the authors, who reviewed and approved the final version of the manuscript. AI output was revised and modified by the authors before submission.

## Acknowledgements

FSB did all the wet lab experiments, did the Epi2ME analysis, help drafted the manuscript and revised the final version. WPT developed the sequencing protocols, sequenced all samples, wrote the relevant methodology section and revised the final manuscript. AT provided funding, created the EMU workflow and did the bioinformatic, formatted and analyzed the results, drafted the article and revised all versions.

This work was supported by Canada NSERC discovery program, grant number RGPIN-2024-04689.

